# Pubertal Testosterone Correlates with Adolescent Impatience and Dorsal Striatal Activity

**DOI:** 10.1101/543710

**Authors:** Corinna Laube, Robert Lorenz, Wouter van den Bos

**Affiliations:** Center for Adaptive Rationality, Max Planck Institute for Human Development, Berlin, Germany

**Keywords:** Testosterone, puberty, impulsivity, intertemporal choice, adolescence

## Abstract

Recent self-report and behavioral studies have demonstrated that pubertal testosterone is related to an increase in risky and impulsive behavior. Yet, the mechanisms underlying such a relationship are poorly understood. Findings from both human and rodent studies point towards distinct striatal pathways including the ventral and dorsal striatum as key target regions for pubertal hormones. In this study we investigated task-related impatience of boys between 10 and 15 years of age (*N* = 75), using an intertemporal choice task combined with measures of functional magnetic resonance imaging and hormonal assessment. Increased levels of testosterone were associated with a greater response bias towards choosing the smaller sooner option. Furthermore, our results show that testosterone specifically modulates the dorsal, not ventral, striatal pathway. These results provide novel insights into our understanding of adolescent impulsive and risky behaviors and how pubertal hormones are related to neural processes.

## 1 Introduction

A recent study reported that adolescents from around the globe (11 countries on four continents, *N* = 5,000) show elevated levels of sensation seeking and immature self-regulation (Steinberg et al., 2017). While increased impulsivity in adolescence may have an adaptive function (Spear, 2000), it may also contribute to significant increases in deaths and accidents during this specific developmental phase (Dahl, 2004). However, the developmental mechanisms driving impulsivity are still poorly understood. Recent research has emphasized *puberty* as a key maturational process involved in impulsive behavior (Crone & Dahl, 2012; Forbes & Dahl, 2010; Laube, Suleiman, Johnson, Dahl, & van den Bos, 2017; for a review see also Laube & van den Bos, 2016). Puberty is defined as the onset of adolescence and is characterized by a surge in pubertal hormones, including testosterone. These hormones are not only involved in changes in the body but are hypothesized to also impact brain functioning. The modulation of specific brain regions is thought to lead to changes in behavior promoting impulsivity and risk taking (Blakemore, Burnett, & Dahl, 2010; Crone & Dahl, 2012; Forbes & Dahl, 2010; Peper & Dahl, 2013). Yet to date, little is known about how pubertal testosterone impacts the neural processes involved in decision-making. In the current study we sought to elucidate possible mechanisms underlying impulsive behavior in adolescence that are related to the increase in pubertal testosterone.

Studies on the relationship between pubertal hormones and risky and impulsive behavior in adolescence have focused on testosterone in both boys and girls. During puberty, testosterone levels rise for both sexes, yet this increase is steeper and more prolonged for males (Braams, van Duijvenvoorde, Peper, & Crone, 2015). Indeed, several studies have reported a positive relationship between pubertal testosterone and self-reported (Vermeersch, T’Sjoen, Kaufman, & Vincke, 2008) and observed (Cardoos et al., 2017; Forbes et al., 2010; Op De MacKs et al., 2011) risk-taking. Furthermore, we recently found a similar positive relationship for pubertal testosterone and impatience (Laube et al., 2017), an important component of impulsive behavior (Romer, 2010). Specifically, we found that in early-adolescent boys (11-14 years), increased levels of pubertal testosterone were associated with an increased sensitivity to immediate rewards, whereas age was negatively correlated with the extent to which future rewards were discounted (Laube et al., 2017). In sum, observational and self-report studies support the hypothesis that pubertal testosterone is related to increases in impulsive behavior. Recent neuroscientific findings suggest two possible mechanisms for these increases.

First, pubertal testosterone may specifically impact computations in the ventral striatum. It is well known that the ventral striatum plays a key role in representing subjective value in decision making (Bartra, McGuire, & Kable, 2013; Haber & Knutson, 2009; Moreira, Pinto, Almeida, Barros, & Barbosa, 2016). More importantly, there have been several imaging studies in humans that also point towards a positive relationship between pubertal testosterone and the ventral striatum (for a review see Laube & van den Bos, 2016). For example, a recent longitudinal study by Braams and colleagues (2015) with a large sample (*N* = 299) found that the developmental trajectory of nucleus accumbens (NAcc) activity related to receiving probabilistic rewards showed a peak in activation during adolescence. Moreover, NAcc activation was positively correlated with salivary testosterone. In line with these findings, Op de Macks and colleagues (2011) found that testosterone levels were positively correlated with activation within the NAcc in response to rewards. Taken together, and in line with most dual-process theories of adolescent decision making (Casey, 2014; Luna, Marek, Larsen, Tervo-Clemmens, & Chahal, 2015; Shulman et al., 2016; Steinberg, 2010), these findings support the hypothesis that pubertal testosterone is modulating *valuation processes* in the ventral striatum, which may lead to different representations of immediate rewards and more impulsive behavior.

Second, human and animal studies have identified the dorsal striatum as a key target of pubertal hormones (Matthews, Bondi, Torres, & Moghaddam, 2013; Sinclair, Purves-Tyson, Allen, & Weickert, 2014; Laube & van den Bos, 2016). More specifically, animal studies have suggested that there is a decrease in dopamine function in the dorsal striatum due to a rise in pubertal testosterone in male adolescents (Matthews et al., 2013; Purves-Tyson et al., 2014; Stamford, 1989). For instance, Purves-Tyson and colleagues (2014) found that dopamine activity in the dorsal striatum of mice was increased after gonadectomy and weakened by testosterone replacement. Furthermore, Matthews and colleagues (2013) found that diminished presynaptic dopamine activity in male adolescent mice was limited to the dorsal, but not the ventral, striatum. Both the ventral and the dorsal striatum receive projections from the dopamine system (Haber & Knutson, 2009), yet from distinct cortical areas (Tziortzi et al., 2014). The ventral striatum receives afferent projections from areas associated with reward processing, including limbic regions, while the dorsal striatum receives its main afferent connections from the frontal cortex, an area associated with executive control (Balleine, Delgado, & Hikosaka, 2007; Haber & Knutson, 2009; Tziortzi et al., 2014). In line with its connectivity profile, the dorsal striatum is associated with top-down modulation of learning and action selection (Dayan & Berridge, 2014; Frank, 2011). For instance, we previously found that the strength of the structural connectivity between the frontal cortex and the dorsal striatum is associated with increased future-orientation and reduction of impulsive decision making (van den Bos, Rodriguez, Schweitzer, & McClure, 2014; van den Bos, Rodriguez, Schweitzer, & McClure, 2015). Taken together, these results indicate that pubertal increases in testosterone may modulate processes in the dorsal striatum, specifically via reducing the dopamine availability. This altered dopamine activation may lead directly to a *bias* towards more impatient behavior (Smith et al., 2016; see also Rigoli, Friston, & Dolan, 2016; Rutledge, Skandali, Dayan, & Dolan, 2015 for relation with dopamine modulation and increased response bias).

In sum, testosterone’s distinct effects on ventral and dorsal striatal regions may represent two pathways for influencing decision-making processes in early adolescence. More specifically, the literature suggests that testosterone may modulate dopamine related activity in (1) the ventral striatum which affects valuation related activity, or (2) *modulate* the dorsal striatum and bias action selection. Clearly, these processes are not mutually exclusive and may operate in parallel. Indeed, we have recently shown that different corticostriatal circuits have opposing, and dissociable, effects on impulsive behavior (van den Bos et al., 2014). In a follow-up study we showed that age-related changes in adolescent impulsivity were associated with immature dorsal striatal circuitry, but pubertal hormones were not assessed (van den Bos et al., 2015).

The current study was designed to test these alternative hypotheses about how pubertal testosterone may affect impulsive behavior in adolescence via distinct striatal pathways. To this end we investigated developmental differences in impatience of *N* = 75 boys between 10 and 15 years of age, using an intertemporal choice task. The task, partly performed in a magnetic resonance imaging (MRI) scanner, was designed to specifically test for sensitivity to immediate rewards. To measure pubertal testosterone, we collected two independent morning saliva samples (cf. Laube et al., 2017). The focus of our imaging analyses was on ventral and dorsal striatal regions of interest (ROIs) that were selected on the basis of their connectivity patterns (Tziortzi et al., 2014; van den Bos et al., 2015). Finally, we used computational modeling to gain further insight into the specific cognitive processes that underlie puberty-related changes in impulsivity, specifically aimed at disentangling the processes related to value calculation (ventral striatum) and action selection (dorsal striatum). This multimodel, multimethod approach provides novel insights into the role of pubertal testosterone in brain function and behavior in adolescence.

## Methods

### Participants

As a direct follow-up on to our previous article (Laube et al., 2017) our study focused only on adolescent boys. Adolescent boys (*N* = 75) between the ages of 10 and 15 years (*M* = 12.56 years, *SD* = 1.64) were recruited via the participant database of the Center for Adaptive Rationality at the Max Planck Institute for Human Development in Berlin, Germany and were screened for MRI exclusion criteria (e.g., non-removable ferromagnetic material). Included were boys who were currently enrolled in school, medically healthy with no history of neurological or psychiatric illness, native German speakers, and free from contraindications to MRI. Furthermore, there were no age-related differences in cognitive functioning, as measured by their performance on the Culture Fair Intelligence Test (see Supplementary Materials Table S1).

Before the boys participated in the study, written informed consent was obtained from their parent or legal guardian. Seventy-five participants completed the entire study. These participants received 60 euros.

Five participants were excluded from analysis for the following reasons: (1) task-related imaging data were invalid due to movement (*n* = 3; see also section on functional MRI [fMRI] preprocessing for specific criteria); and (2) data collection was canceled because participants communicated feelings of discomfort and fear inside the MRI scanner (*n* = 2). In total, we had *N* = 70 participants for the behavioral analysis.

### Pubertal Measures

#### Testosterone

Testosterone levels were measured via two morning saliva samples provided by each participant, which is a well-validated method for assessing general circulation of testosterone (Laube et al., 2017; Shirtcliff, Dahl, & Pollak, 2009). We used the passive drool method of saliva collection to minimize discomfort and maximize compliance. Participants were instructed to collect the two saliva samples on separate—preferably consecutive—mornings, ideally 15–30 min after waking, and to immediately place the samples in the freezer. They completed a form indicating the date and time each sample was collected. Participants came into the laboratory twice: At their first visit, they were given the information and two empty devices for saliva collection, which they were told to return upon arrival for their second visit. Importantly, the time between the first and second visit was a maximum of 7 days. When brought to the lab, the saliva samples were immediately stored in a freezer at −20 °C. Subsequently, they were frozen at −80 °C for long-term storage. Testosterone assays were conducted at ISD Laboratory. The intra-assay correlation was high, *r*(68) = .73, 95% confidence interval (CI) [.61, .82], *p* < .001. Testosterone levels were therefore calculated as the average across the two samples collected by each participant. In one case, one of the two samples was excluded because it was too contaminated to be analyzed. In this case the value of the valid sample was used, rather than the average. See Figure S1 and Table S1 in Supplementary Materials for more information. Note that testosterone levels were non-normally distributed (Shapiro–Wilk’s *W* = 0.91, *p* < .001) and were log transformed for further analysis.

#### Self-reported pubertal stage

We also administered the Pubertal Development Scale (PDS) self-report measure to estimate pubertal stage (Petersen, Crockett, Richards, & Boxer, 1988). It is commonly used to assess *external* pubertal status and asks adolescents about hair growth, skin changes, and growth spurts, resulting in a composite puberty score. Although the PDS scale is a less specific measure of pubertal development than hormonal assessment, it was included as an additional measure of pubertal status because it captures the physical changes triggered by testosterone. As expected, the PDS score and pubertal testosterone were positively correlated, *r*_s_ = .76, 95% CI [.64, .84], *p* < .001. Seventeen subjects scored 1 on the PDS but had testosterone levels > 11 pmol/L, while one subject had a testosterone level of 5.7 pmol/L but a PDS score of 1.3. From this we can conclude that every subject (*N* = 70) in the current study had started puberty (see Table S1 in Supplementary Materials for more information).

### fMRI Paradigm

Participants made 64 binary choices between two hypothetical amounts of money available at different delays (see Figure 1). The smaller–sooner (SS) option offered a small reward after a short delay; the larger–later (LL) option offered a larger reward after a longer delay. For half of the trials, the smaller reward was available “today”; for the other half, both the smaller and the larger reward were available in the future, at different delays. Importantly, the stimulus set used in the scanner was specifically tailored to the temporal preferences of each individual participant (see also Rodriguez, Turner, Van Zandt, & McClure, 2015; van den Bos et al., 2015). That is, to achieve an equal distribution of SS and LL choices and create a stimulus set that was comparable across participants in terms of differences in subjective value, we added an additional adaptive intertemporal choice task prior to the fMRI task. This task consisted of a total of 60 trials, which were determined by a staircase procedure. For this, the SS reward was fixed at 10 euros received today. The delay period for the LL reward was randomly chosen from a uniform distribution between 15 and 60 days in the future. The size of the LL reward was adjusted to converge toward the same subjective value as the SS outcome. After the participants completed 60 trials of this task, we fitted data with the hyperbolic discount function whose parameter estimates were used to generate the trials of the scanner session (see Supplementary Materials for details). Importantly, neither the monetary amounts nor the generated time delays for the fMRI task correlated with testosterone, age, or PDS (all *p* > .05).

**Figure 1.**
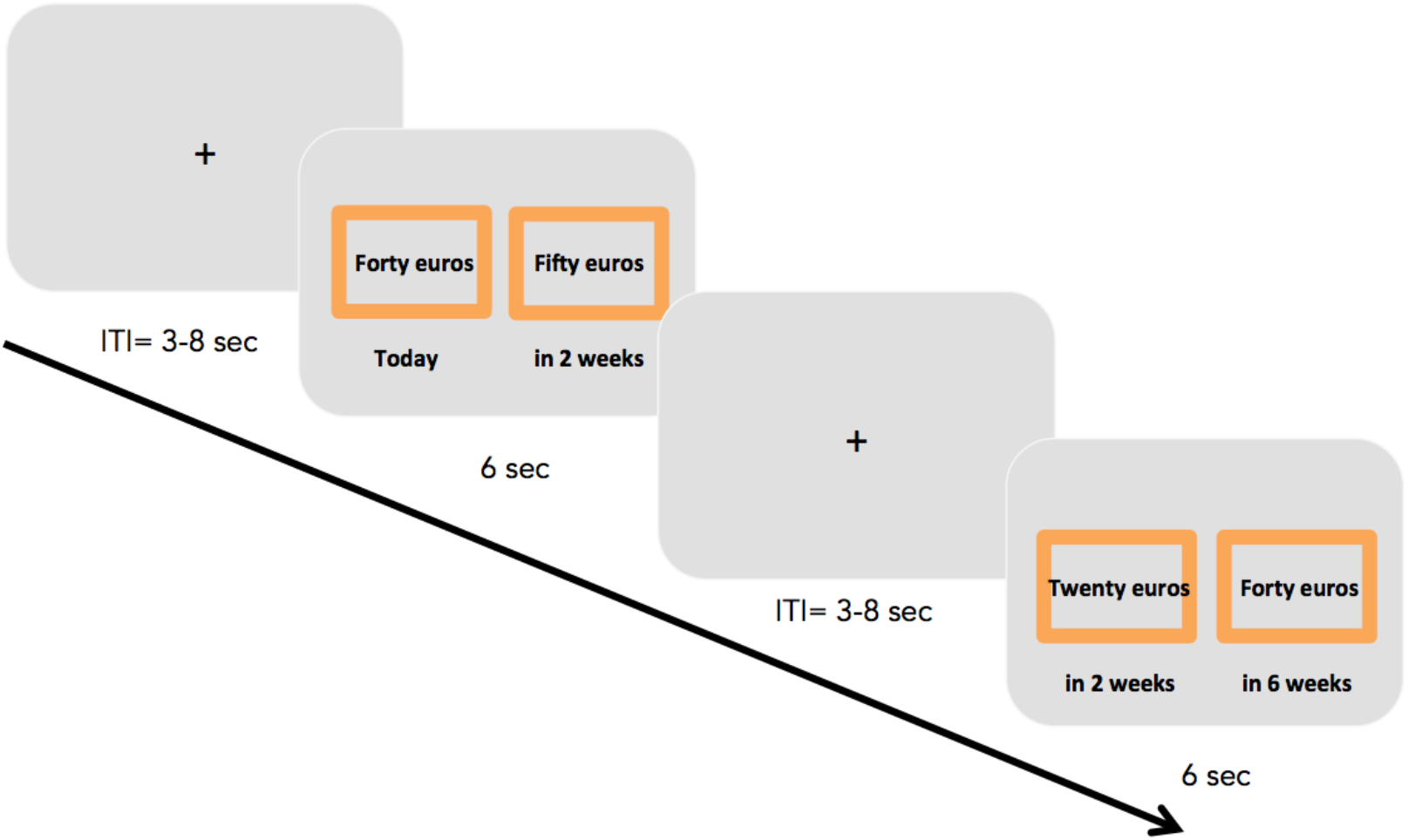
Example of the paradigm used in the current study. ITI = intertrial interval.

Although participants were not directly paid the actual monetary amounts used in the task, past research has consistently shown that choices with hypothetical and real rewards in a delay-discounting paradigm significantly correlate with each other (Bickel, Pitcock, Yi, & Angtuaco, 2009; van den Bos et al., 2015). In addition, evidence on incentives used in intertemporal choice studies has demonstrated small effects on choice behavior (Augenblick, Niederle, & Sprenger, 2015).

The inter-trial interval was 2-8 seconds (*M* = 4.5 seconds). Each choice was presented for 6 seconds, the maximum amount of time participants had to indicate their answer (see Figure 1).

### Behavioral Analyses

To gain insight into the underlying decision strategies, we modeled participants’ choices using three types of discounted utility models. The basic assumption of these models is that when a reward is available at a certain delay, its subjective value is discounted relative to the extent of that delay:

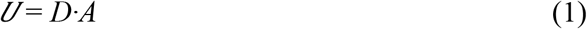

where *U* is the subjective utility and *A* is the objective monetary amount, which is multiplied by a discount function *D*. According to the canonical utility models, all one needs to understand is how to characterize *D*. In the classic hyperbolic discounting function, *D* is defined as

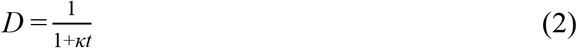

where *t* is time and *κ* is the discount factor (greater □ implies greater impulsivity).

To better capture individual differences in sensitivity to immediate rewards as opposed to long-term rewards, we also used two well-known two-parameter discount models:

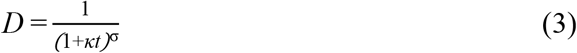

In this two-parameter model σ reflects individual differences in sensitivity to change at shorter delays relative to longer delays. Given its increased flexibility, the two-parameter model often shows increased fit and predictive power, compared to the single-parameter hyperbolic model (Green & Myerson, 2004; Laube et al., 2017; Wulff & van den Bos, 2016). In our previous research, we found that this model best described the behavior of an adolescent population, and importantly that pubertal testosterone was specifically associated with the *σ* parameter, indicating that testosterone is associated with increased behavioral sensitivity to near-term rewards. Here we extend our previous modeling efforts to further identify the possible mechanism that underlie impatience. First we focus on another type of two-parameter model that suggests there may be multiple systems that discount rewards at different rates (McClure et al., 2004, 2007; Laibson, 1997; Radu et al., 2011). Here we test the simple version of such a model based on its relatively good performance in a larger model comparison set using a big data set (Wulff & van den Bos, 2017). This model assumes that there are two systems that apply different discount rates:

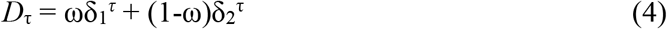

where ω indicates the relative involvement of each system in a given decision. Here, exponential discount rates δ_1_ and δ_2_ must be at least 0 and no greater than 1 in value. Second, δ_1_ < δ_2_, given that we assume that there is one system discount that is greater than the other. Finally, the weighting parameter ω must be between 0 and 1. The model assumes that the more impatient system, δ_1_, is associated with the ventral striatum and connected limbic regions. On the basis of our earlier findings we would therefore expect that higher levels of pubertal testosterone would be associated with either a higher discount rate for δ_1_ or a stronger involvement (higher ω) of the δ_1_ system in behavior. Note that although they have similar effects on impatience they represent different psychological mechanisms: One impacts the valuation process directly, and the other more-or-less biases choice without directly changing the valuation of the rewards. As such, they map onto the proposed ventral (valuation) versus dorsal (action selection/executive control) striatum distinction. For all of the above models we used the logistic choice rule to compute the probability of choosing the SS option as a function of the difference in subjective values *V*_SS_ and *V*_LL_:

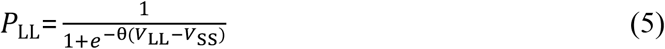

where θ estimates response noise. This function assumed that each individual would choose the option with the highest subjective value with the highest probability.

To further explore the distinction between processes associated with the ventral and dorsal striatum, we included two more models that could potentially capture the distinction between evaluative and non-evaluative selection processes. In contrast to the dual-system models, these models assume that there is a single evaluation system (Kable & Glimcher, 2007; Rangel, Camerer, & Montague, 2008) that discounts future rewards hyperbolically, as described by the canonical hyperbolic model (see Equation 2). In addition, these models include a response-bias parameter inspired by the response bias that is usually implemented in sequential sampling models of choice (Ratcliff & Smith, 2004; see also Rodriguez 2014, 2015 for such a model implementation in the context of intertemporal choice). Implementation of the bias parameter should be regarded as a pre-evaluative process, such as setting an aspiration level of receiving something now, as soon as possible, or sometime in the future (Ericson et al., 2015; Stewart, Reimers, & Harris, 2014; Wulff, Hills, & Hertwig, 2015; Wulff & van den Bos, 2017). We tested two different bias models: (1) an immediacy bias model (IMM_bias_) and (2) an as-soon-as-possible bias model (SS_bias_). Following Stewart and colleagues (2014; see also *Rutledge et al, 2015; Rigoli et al, 2016*), these models operate on the choice function itself:

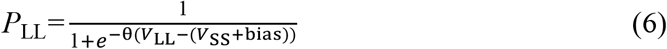

Here the bias parameter allows for an overall bias toward SS or LL choices independent of the amounts and delays associated with the options. A positive bias indicates a bias toward SS options and a negative value indicates a bias toward LL options. For the IMM_bias_ model the bias parameter is 0 when the SS option is also in the future and a free parameter when the SS option is immediate. We used the optimization toolbox optim implemented in R for model fitting (Nash & Varadhan, 2011). Maximum likelihood of the observed data was calculated with the choice functions in Equations 5 and 6. The Bayesian information criterion provided an indication of the relative fit of the statistical models given the data; cross-validation using the logLoss scoring rule was used to gauge the relative differences in predictive accuracy (see Supplementary Materials for details). The models showed distinct differences in fits to the choice data, but their levels of out-of-sample predictive accuracy were similar.

### MRI Image Acquisition

MRI scans were conducted with a 3 Tesla Siemens TIM Trio Scanner (Erlangen, Germany) at the Max Planck Institute for Human Development in Berlin, Germany. The intertemporal choice task was presented via a video projector and a mirror system mounted on top of the 12-channel phased array head coil. We collected 346 functional images per participant using a T2*-weighted echo planar imaging (EPI) sequence that was individually aligned to the genu and the splenium of the corpus callosum. The following parameters were used: 36 slices, ascending interleaved slice order, time to repeat (TR) = 2 s, time to echo (TE) = 30 ms, field of view (FoV) = 216 × 216, flip angle = 80°, matrix size = 72 × 72, voxel size = 3 × 3 × 3 mm^3^. Additionally, three-dimensional anatomical images of the whole brain were obtained with a T1-weighted magnetization-prepared rapid gradient echo (MP-RAGE) sequence to optimize the normalization procedure during data preprocessing (TR = 2.500 ms, TE = 4.77 ms, inversion time = 1,100 ms, acquisition matrix = 256 × 256 × 192, flip angle = 7°, voxel size: 1 × 1 × 1 mm^3^).

### fMRI Preprocessing

Analysis of the fMRI data was conducted using Statistical Parametric Mapping 8 (SPM8, Wellcome Department of Imaging Neuroscience, London, UK). Functional images were corrected for acquisition time delay and head motion. Then, the anatomical reference image was coregistered to the mean EPI image and transformed into the stereotactic normalized standard space of the Montreal Neuroimaging Institute (MNI) using the unified segmentation algorithm as implemented in SPM8. Then, EPIs were spatially normalized (using the normalization parameters computed during segmentation), resampled (voxel size = 3 × 3 × 3 mm^3^), and spatially smoothed with a three-dimensional Gaussian kernel of 6 mm full width at half maximum. For display purposes, a mean anatomical image of the sample was computed from the spatially normalized individual images.

Finally, we used ArtRepair, a toolbox for SPM (Mazaika et al., 2009), to identify outlier scans caused by large sharp movements (cut-off value of 3 *SD*) and by global mean intensity drifts (cut-off value of 2 SD). Using this procedure an average of 9.24 scans (*SD* = 15.89, range: 0–60) were identified (in all participants below 20% of 346 scans total). Participants were excluded from analysis if more than 20% of their volumes had to be removed (*n* = 3; see Participants, above).

### fMRI Analyses

Imaging data were analyzed using a two-stage mixed-effects general linear model. A subject-specific design matrix was created to test for brain responses associated with impatience (choice model). The choice model included two regressors of interest that modeled SS and LL choices for every trial. Each regressor was convolved with the hemodynamic response function provided by SPM8. Additionally, the six rigid body-movement parameters, mean signal drift, and the outlier scans (for each outlier scan a regressor containing a 1 for the outlier scan and 0 for all others was created) as identified by the ArtRepair toolbox (see fMRI Preprocessing, above) were included as regressors of no interest.

The current study was designed to test the relationship between testosterone and the sub-regions within the striatum, and how these relationships are reflected in impatient choice. To test this, we performed ROI analyses using the MarsBaR toolbox (Brett et al., 2002; http://marsbar.sourceforge.net/) for SPM8.

Importantly, in recent research, the functional subdivision of the striatum was based on distinct afferent projections (providing input) from cortical areas (Tziortzi et al., 2014). While the ventral striatum receives input mainly from regions associated with different aspects of reward and emotional processing, such as the ventromedial prefrontal cortex, orbitofrontal cortex, and dorsal anterior cingulum, as well as limbic regions such as the amygdala and hippocampus, the dorsal striatum receives converging projections from areas involved in executive functioning, such as the medial prefrontal cortex (Balleine et al., 2007) and the dorsolateral prefrontal cortex, as well as regions involved in motor control and visual attention, such as the supplementary motor area and the frontal eye field region, respectively (Tziortzi et al., 2014). Thus, to precisely capture the underlying functions associated with different areas within the striatum, ROI analyses on two a priori selected striatal regions were based on their probabilistic connectivity according to cortical–striatal anatomical connections with three cortical targets: limbic, executive, and rostral motor. ROIs were derived from the seven sub-regions defined by the Oxford–GSK–Imanova Striatal Connectivity Atlas (Tziortzi et al., 2014); see Figure 2 for ROIs).

**Figure 2.**
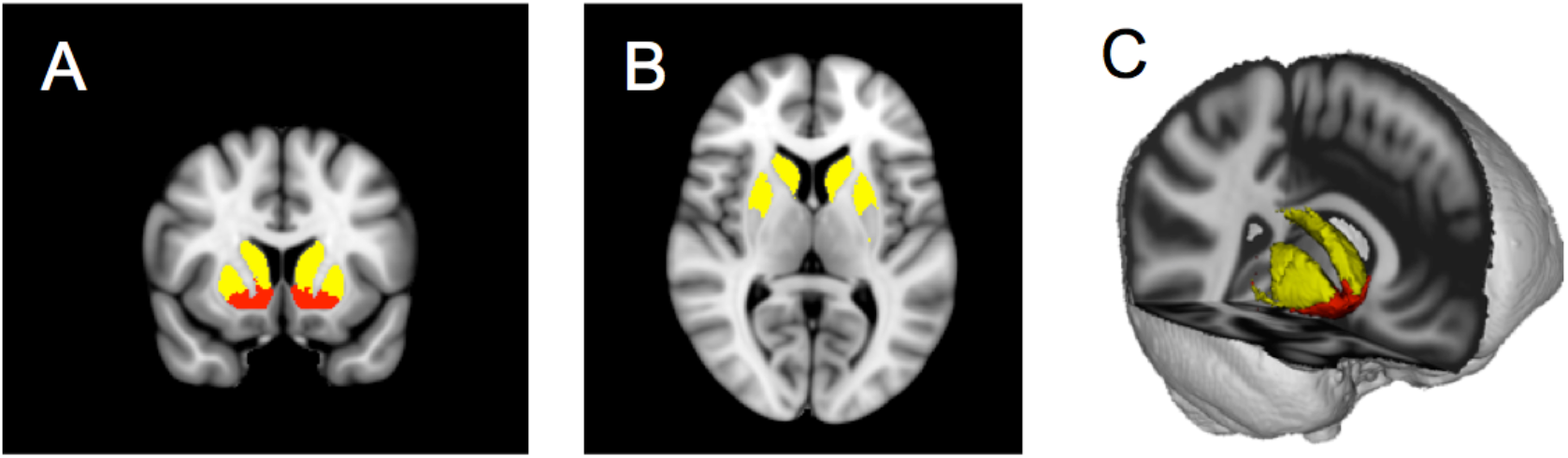
Bilateral regions of interest (ROIs) used in all imaging analyses were based on functional masks (Oxford–GSK–Imanova Striatal Connectivity Atlas). (A) Nucleus accumbens (red), dorsal striatum (yellow). (B) Dorsal striatum axial view. (C) Three-dimensional view of both ROIs with ventral striatum (red) and dorsal striatum (yellow).

We performed ROI analysis and used these ROIs for small-volume correction. Given our ROI focused approach we used a stricter threshold than is normally used on the whole brain level. That is, the standard familywise error (FWE) threshold of .05 was again Bonferroni-corrected to .025 in order to consider task-related responses as significant. Results remain similar with a more liberal threshold.

First, we were interested in how these two distinct striatal parts were involved in impatient decision making. Therefore, we contrasted activation associated with the SS over LL choices to investigate how ventral and dorsal striatal regions were involved in either impatient or patient choices. In a subsequent step, we examined how levels of pubertal testosterone were associated with the choice-related activity in the ventral and dorsal striatum. Because we were interested in the unique contribution of pubertal testosterone to brain activity we controlled for age in all of our analyses (similar to our behavioral analyses). Even though our procedure with the pre scanner task was aimed at getting the ratio of SS and LL choices close to 50–50 for each participant, there were still some individuals who significantly deviated from this distribution. To reliably estimate the SS–LL contrast we included only individuals who made at least 20% of SS or LL choices, resulting in a sample of *N* = 48 (see Table S4 in the Supplementary Materials for the summary statistics for this imaging group).

Second, even though we did not find any behavioral immediacy effect, we did compare now–later with later–later options to see if the ventral, and possibly dorsal, striatum was more sensitive to the presence of immediate rewards (controlling for choice type). Similar to in the previous analyses, we subsequently added testosterone as a regressor of interest and age as a regressor of no interest to this contrast (all participants, *N* = 70, were included in these analyses).

To check the robustness of our results we repeated the analyses using all participants (*N* = 70) for the SS–LL contrast, and in the subgroup in which collinearity between age and testosterone was reduced (*N* = 32, see Behavioral Analyses, above) for both contrasts. All of the reported results hold when using these groups (see Supplementary Materials), suggesting that the reported results are robust to individual differences in behavioral patterns and individual differences in age and testosterone.

## Results

### Behavioral Results

#### Summary statistics and regression analysis

As expected, the correlation with testosterone levels was strong for self-reported pubertal development (PDS), *r* = .76, 95% CI [.73, .79], *p* < .001, which supports the idea that individual differences in testosterone levels are related to pubertal development. Furthermore, there was a correlation between testosterone and age, *r* = .73, 95% CI [.70, .76], *p* < .001 and age and PDS, *r* = .80, 95% CI = [.78, .82], *p* < .001 (see also Table 1 for all zero-order correlations between all study variables). Finally, the overall distribution of SS and LL choices was relatively equal (*M*_SS_= 56%, *SE* = 2.84%), which we also aimed to achieve with the adaptive design.

**Table 1.** Means and Standard Deviations for the Study Variables, and Zero-Order Correlations With 95% Confidence Intervals

| Variable | Age | Testosterone | PDS | *M* | *SD* |
| --- | --- | --- | --- | --- | --- |
| Age (in months) | — |  |  | 156.99 | 19.11 |
| Testosterone | .73*** | — |  | 78.29 | 53.90 |
| (pmol/L) | [.70, .76] |  |  |  |  |
| PDS | .80*** | .76*** | — | 1.82 | 0.71 |
|  | [.78, .82] | [.73, .79] |  |  |  |

*Note.* $N = 70$ . PDS = Pubertal Developmental Scale.
\*\*\* $p < .001$ .

#### Bias and testosterone

Bayesian model comparison indicated that the SS bias model fit the data best (see Table 2 for model fits and Table 3 for the best fitting parameters). Since we were interested in the unique contribution of testosterone in impatient choice, we performed a multiple regression analysis to predict the level of bias for SS options based on both testosterone and age. Testosterone, but not age, was a significant predictor for the *bias* parameter, *b* = .38, 95% CI [.04, .73], *p* = .03 (see Figure 3), and *b* = .14, 95% CI [−.66, .05], *p* = .10, respectively. Thus, those adolescents who had higher levels of testosterone also showed an increased response bias to the SS option. However, there was no significant effect of testosterone on the discount parameter *k, b* = .05, 95% CI [−.07, .47], *p* = .81. Similarly, age also did not have a significant effect on *k, b* = −.30, 95% CI [−.21, .48], *p* = .43.

**Table 2.**
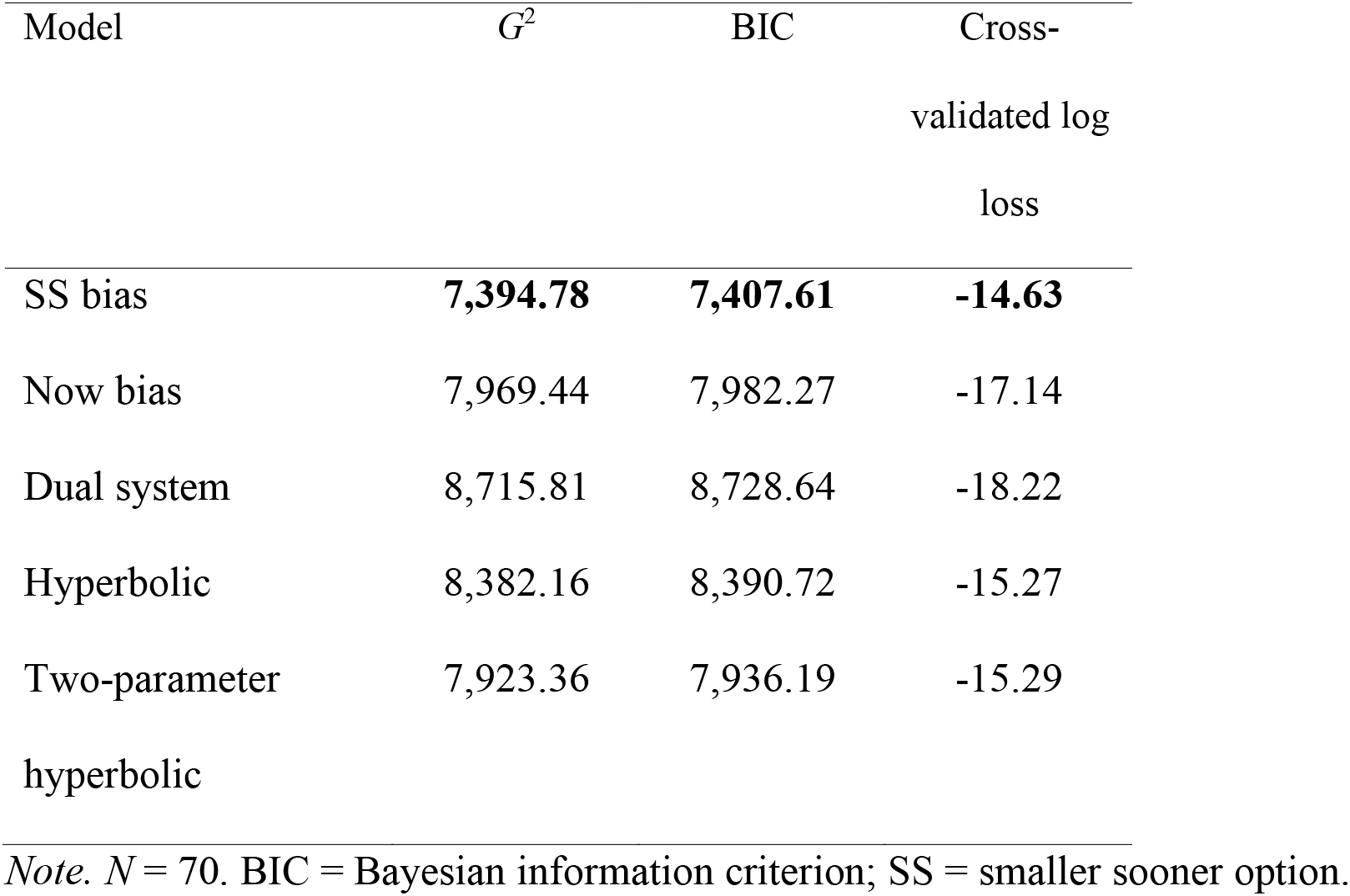
Model Fits for Different Discounting Models

| Model | $G^2$ | BIC | Cross-<br>validated log<br>loss |
| --- | --- | --- | --- |
| SS bias | **7,394.78** | **7,407.61** | **-14.63** |
| Now bias | 7,969.44 | 7,982.27 | -17.14 |
| Dual system | 8,715.81 | 8,728.64 | -18.22 |
| Hyperbolic | 8,382.16 | 8,390.72 | -15.27 |
| Two-parameter<br>hyperbolic | 7,923.36 | 7,936.19 | -15.29 |

*Note.* $N = 70$ . BIC = Bayesian information criterion; SS = smaller sooner option.

**Table 3.** Mean Parameter Estimates SS Bias

| SS bonus |  |
| --- | --- |
| $\kappa$ | .08 (.003) |
| Bias | 14.29 (2.80) |

*Note.* $N = 70$ . None of the parameter estimates were correlated with each other (all $ps$ $> .2$ and all $rs < .15$ ). SS = Smaller sooner option.

#### Outliers

Because there were some outliers within the distribution of the bias parameter, we used the interquartile range method to remove these (minimum value: −38, maximum value: 59) and repeated our previous analyses. The significant relationship between testosterone and bias still remained, *b* = 0.39, 95% CI [0.03, .0.75], *p* = .04, whereas age again did not show a significant relationship with bias, *b* = −.30, 95% CI [-0.66, .05], *p* = .04. Confirming the previous results, there was again no significant relationship between *k* and testosterone, *b* = 1.41, 95% CI [−.66, .05], *p* = .13, or age, *b* = −.01, 95% CI [−.66, .05], *p* = .61.

#### Testosterone and Age

Next, because of the positive correlation between testosterone and age (*R*^2^ = .53), we also created a subsample in which the correlation was lowered (*R*^2^ = .26; see Supplementary Materials Table S3 for more summary statistics). Here, we calculated quartiles based on age and used the two middle quartiles for the subsample, which resulted in a sample with *N* = 32 participants. While the relationship between testosterone and age was still significant (*p* < .01), the variance inflation factor was 1.35, as opposed to 2.09 for the entire sample, which indicates that the shared variance (or multicollinearity) between age and testosterone regressors was not problematic for fitting the model. As expected, the relationship between the bias parameter and testosterone remained significant in a multiple linear regression controlling for age, *b* = .61, 95% CI [.23, .99], *p* = .003. Again, a significant effect of testosterone on the discount parameter *k* was not apparent, *b* = .04, 95% CI [−.40, .48], *p* = .85.

Taken together, these results are in line with our previous findings(Laube et al., 2017), as they indicate that higher levels of testosterone are specifically related to a bias for choosing impulsively.. That is, even if the subjective values of the SS and LL rewards were the same, participants would still be more likely to choose the SS option. Alternatively, they would only choose the LL option if its relative subjective value was higher than the bias. In sum, results suggest that testosterone biases choices in a way that is independent of value calculation, which is consistent with its relationship with the dorsal but not the ventral striatal pathway.

### Imaging Results

#### Choice-related activity (SS vs. LL)

First, we tested for distinct patterns of activity when individuals chose LL rewards compared with when they chose SS rewards. This contrast identified the left NAcc, *p* = .01 FWE, MNI: -9 5 -5, *t*(47) = 3.37, as being more active when individuals chose the LL compared to the SS option (see Figure 4). Given that LL choices are often only chosen when the subjective value is higher, to overcome the participants response bias to SS options (see above), this increased level of activity may reflect the increased level of subjective value. This interpretation is in line with the current literature showing that activity in the ventral striatum reflects subjective value (for meta-analysis see Batra et al., 2013; see also Neurosynth map associated with “value”). We did not find any significant results for the opposite contrast.

**Figure 4.**
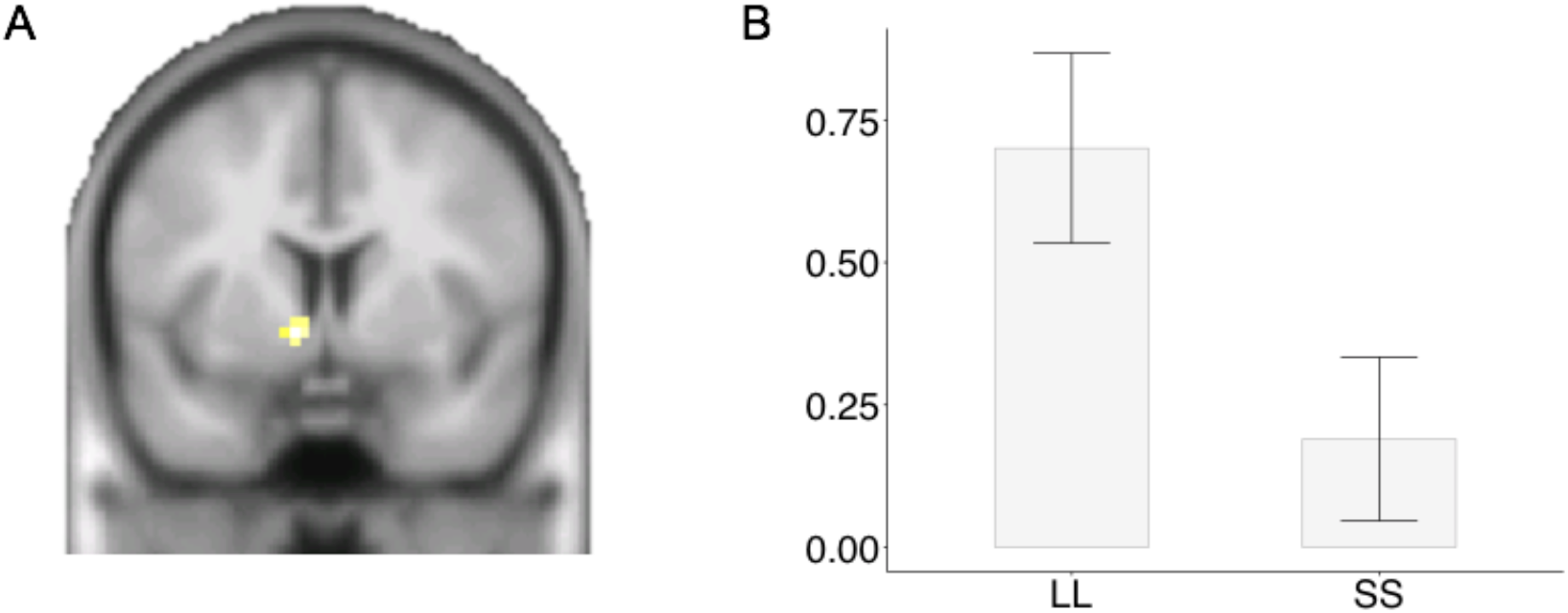
(A & B) Region of interest ROI analysis revealed higher activity in the nucleus accumbens (NAcc) for larger later (LL) compared to smaller sooner (SS) choices.

#### Immediacy effect (now vs later)

Recently, we hypothesized that a relationship between pubertal testosterone and sensitivity to immediate rewards would be modulated by the ventral striatum. In particular, the ventral striatum has been shown to be selectively engaged in immediate choices (McClure, Laibson, Loewenstein, & Cohen, 2004). Thus, even though we did not find a behavioral effect in this sample, we investigated if the context of the choice options is related to activity within the ventral striatum and possibly the dorsal striatum. That is, we compared today–later with later–later options to test if one (or possibly both) of our selected ROIs was more sensitive to the presence of immediate rewards. We included choice as a regressor of no interest, because we were interested in the unique contribution of reward context independent of choice, given that choices were also biased toward an equal distribution between SS and LL choices. However, in line with our behavioral findings, none of these analyses revealed significant results.

### Testosterone and choice

To test for testosterone-specific modulation of neural activation, we performed a regression analysis with testosterone level as a predictor on the SS–LL contrast, again this test was performed for both dorsal and ventral striautm. This analysis resulted in significant activation within the dorsal striatum (caudate nucleus, *p* = .021 FWE, MNI: -12 17 10, *t*(45) = 3.28; see Figure 6). As such, testosterone level predicted the extent of activation in the dorsal striatum, such that higher levels of testosterone corresponded with increased activation for LL compared to SS choices. We did not find any activity associated with testosterone in the ventral striatum (even at more liberal thresholds), nor did exploratory whole brain analyses result in any testosterone sensitive regions.

**Figure 6.**
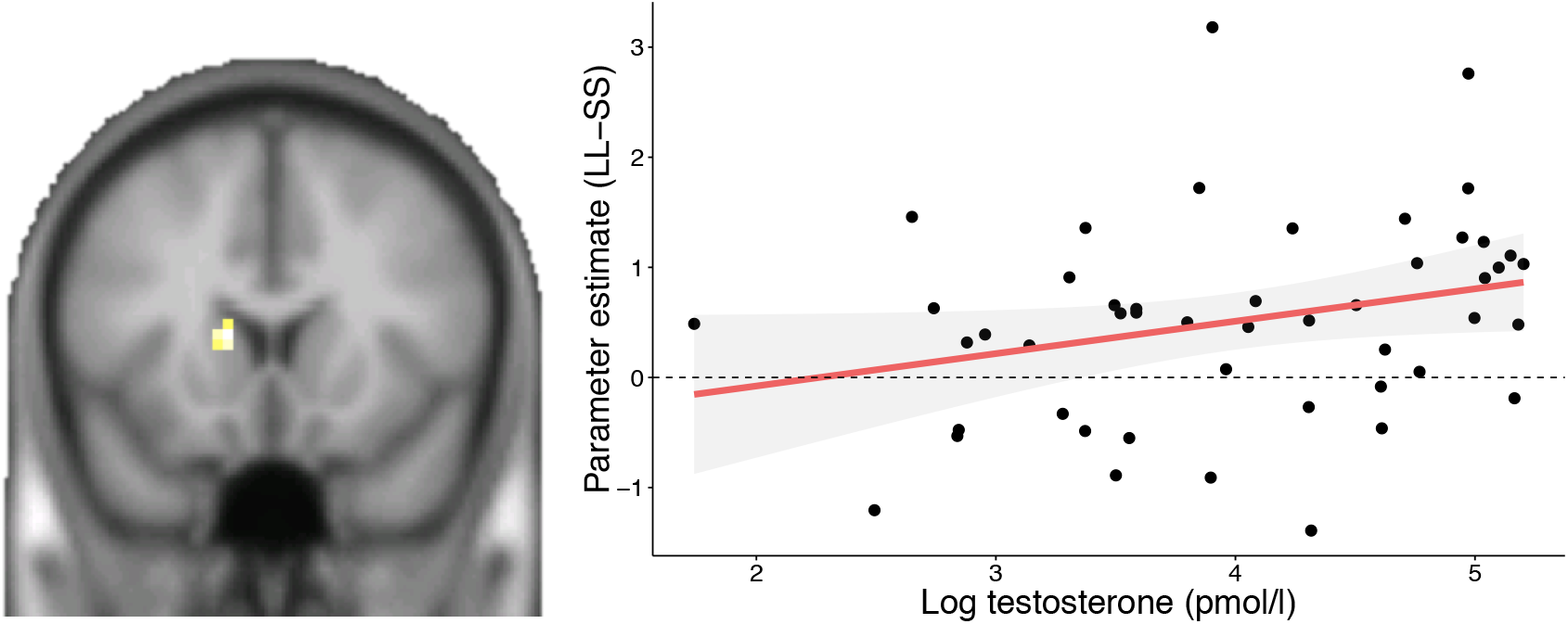
Region of interest (ROI) regression with testosterone as covariate of interest in the dorsal striatum, controlling for age, for the choice contrast between smaller sooner and larger later rewards revealed significant activity in the dorsal striatum.

In sum, these results suggest a relationship between testosterone and the dorsal but not the ventral striatum, where testosterone seems to modulate activity, which biases individuals’ choices toward the SS option. One way to interpret this finding is that more top-down control was needed for these high-testosterone individuals to choose the LL option. This interpretation is based on the fact that the dorsal striatum receives converging projections from the lateral and anterior prefrontal cortex (Haber & Knutson, 2009). To further explores this hypothesis, we conducted exploratory functional connectivity analysis.

### Exploratory connectivity analyses

To explore whether testosterone in the dorsal striatum may modulate functionally coupling with the prefrontal cortex during the delay discounting task, we assessed functional connectivity of a priori defined ROIs using psychophysiological interaction (PPI). Specifically, we focused on functional connectivity between the dorsal striatum ROI and a ROI including the rostral superior and middle frontal gyri and the dorsolateral prefrontal cortex, as those areas have been identified as cortical input to the dorsal striatum, particularly specialized in executive functions; see Tziortzi et al., 2014).

First, we tested for increased functional connectivity during the decision phase of the task relative to baseline. We extracted the mean BOLD time series from the voxels within the bilateral dorsal striatum ROI related to activation of the choice – baseline contrast. Subsequently, we estimated a GLM for every subject that included the following three regressors in addition to the motion parameters: (1) an interaction between mean BOLD response in the dorsal striatum ROI and the mean centered decision phase regressor convolved with the canonical HRF; (2) a regressor specifying decision phases as an indicator function convolved with the canonical HRF; and (3) the original BOLD eigenvariate from the target area (i.e., the first principal component of time-series from the voxels within the bilateral dorsal striatum ROI). Single-subject contrasts were calculated after estimation of the GLM.

As expected, these analyses revealed a significant increase in connectivity between the dorsal striatum and the prefrontal cortex during the choice *p* = .015 FWE, MNI: 33 50 −8, *t*(34) = 4.04 (see Figure 7).

**Figure 7.**
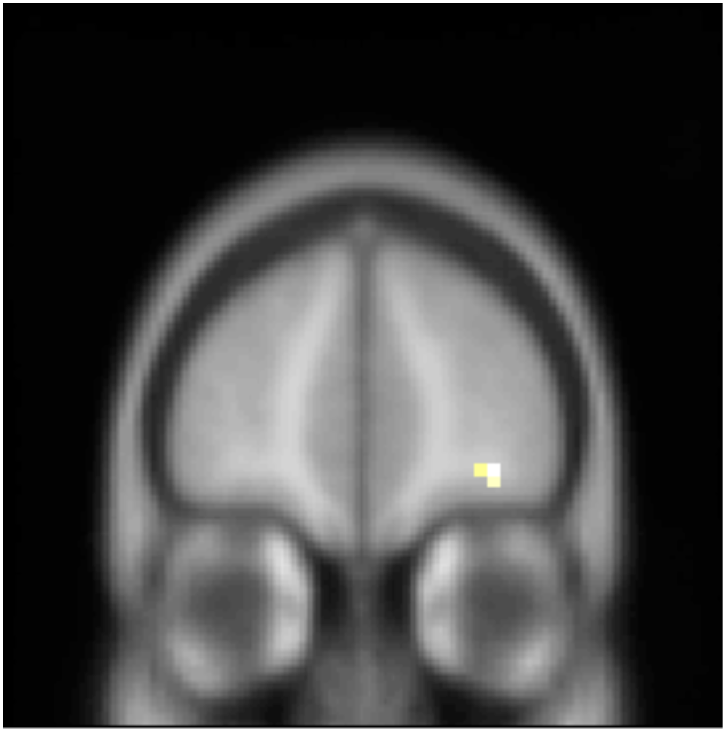
PPI analysis revealed significant functionally connectivity between the dorsal striatum and the prefrontal cortex when making a choice versus baseline.

#### Connectivity & testosterone

Next, we used the Marsbar toolbox to extract mean PPI coefficients from first-level, single-subject contrasts based on the peak-voxel of activation within the bilateral prefrontal ROI that corresponded with the target area used for the PPI analyses. These PPI coefficients were the used as the dependent variable, with testosterone levels as the independent variable, controlling for age. However, we did not find a significant relationship between testosterone level and PPI coefficients, *b* = -0.06, 95% CI [-0.45, 0.33], *p* = .75.

### Brain-behavior

To test the hypothesis that activation within the dorsal stratum is also related to an increase in bias toward choosing the SS option, we conducted a multiple linear regression with beta weights for the SS–LL contrast as the dependent variable, respectively, and the bias parameter obtained by the previous cognitive modeling procedure as the independent variable, controlling for age. As expected we found a negative relationship between the bias parameter and activation within the dorsal striatum for SS over LL choices, *b* = −.18, 95% CI [−.45, .11], *p* = .10, but this failed to reach significance. Consequently, activity in the dorsal striatum may not be directly linked to a behavioral bias for SS options, yet results suggest that there may be an indirect relationship given that the bias parameter may also represent a multitude of various cognitive processes, which eventually may lead to a behavioral bias.

## Discussion

Puberty marks the transition from childhood to adulthood, serving as an important biological marker for tremendous changes in both the structure and function of the brain. However, little is known about how pubertal hormones impact brain function, or how they might affect impulsive behavior in adolescence. The current study was designed to examine the relationship between testosterone and impatient behavior with a specific focus on its interaction with two striatal sub-regions. Our results suggest that testosterone specifically modulates dorsal, not ventral, striatal function in the context of intertemporal choice. Furthermore, increased levels of testosterone were associated with a greater response bias towards choosing the SS option, regardless of its value. These results provide new insights into our understanding of adolescent impulsive and risky behaviors, which we will address below.

Our results are generally consistent with neurocognitive models of delay discounting, which highlight a central role for separate components of the frontostriatal network (Peters & Büchel, 2011; van den Bos & McClure, 2013; van den Bos et al., 2014). First, behavioral results suggest that almost all subjects showed a response bias towards the more immediate option. That is, if the SS and LL options did not differ much in subjective value, people tended to choose the SS option. Thus, to overcome this response bias, the LL option needed to have a higher subjective discounted value than the SS option. Indeed, our imaging results, showing increased ventral striatal activity when participants chose the LL over the SS option, are consistent with numerous previous studies showing that activity in the ventral striatum scales with subjective value (for a review see Bartra et al., 2013).

Second, we found evidence that testosterone modulates dorsal striatal functioning. That is, participants with high levels of testosterone showed relatively more dorsal striatal activation when choosing LL over SS options. Given that the dorsal striatum receives converging projections from the lateral and anterior prefrontal cortex (Haber & Knutson, 2009), this could indicate that more top-down control was needed for these high-testosterone participants to choose the LL option. Indeed, several studies have highlighted a crucial role of prefrontal input to the valuation system in decreasing impatience in both adults (Figner et al., 2010; van den Bos et al., 2014) and adolescents (Achterberg, Peper, van Duijvenvoorde, Mandl, & Crone, 2016; Steinbeis, Haushofer, Fehr, & Singer, 2016; van den Bos et al., 2015).

Furthermore, the increased need for top-down control is consistent with our modeling results showing that higher levels of testosterone were associated with a stronger response bias towards the SS option. One possible mechanism may be that testosterone modulates activity within the dorsal striatum by impacting the dopamine system. Indeed, several rodent studies indicate a link between dopamine functioning and pubertal testosterone in the dorsal striatum (Matthews et al., 2013; Purves-Tyson et al., 2014; Stamford, 1989). More importantly, a recent study by Smith and colleagues (2016) found that an increased preference for immediate rewards was associated with lower dopamine synthesis in the dorsal striatum (Smith et al., 2016). Interestingly, adolescence is also associated with specific changes in dopaminergic signaling (Galvan, 2010; Hartley & Somerville, 2015). Therefore, it would be of great interest to see rodent studies investigating if and how testosterone interacts with adolescence specific changes in dopamine signaling.

Our results speak to imbalance models of adolescent decision making proposing that impulsivity in adolescence results from a combination of an “imbalance” between ventral striatal reward sensitivity and prefrontal cognitive control (Casey, 2014; Ernst, 2014; Shulman et al., 2016; Steinberg, 2010). In contrast to previous developmental studies that focused on outcome processing or anticipation, we did not find a relationship between testosterone and activation within the ventral striatum in relation to choice (Braams et al., 2015; Op de Macks et al., 2011). Here, our findings are in line with previous animal work by Sturman & Moghaddam (2012) reporting differences in neural processing between adolescent and adult rats specifically in the dorsal, but not the ventral striatum. The current results extend our understanding of the relationship between testosterone and striatal functioning in an important way by highlighting the functional difference of sub-regions within the striatum. Instead of proposing a low (Forbes et al., 2010) or high (Braams et al., 2015; Op de Macks et al., 2011) reward sensitivity, we suggest an additional role for testosterone in puberty: biasing action selection. Imaging studies conducted in male adults have already pointed towards testosterone’s modulation of inhibitory control via connectivity between the amygdala and the prefrontal cortex (Peper et al., 2011; Volman, Toni, Verhagen, & Roelofs, 2011). For instance, Volman et al. (2011) showed that lower baseline testosterone levels were associated with more negative functional connectivity between these areas in a social approach-avoidance task. However, here we did not find evidence for altered connectivity, suggesting that testosterone may modulate local control processes within the striatum, most likely through modulating dopamine activity. That is, animal studies have shown that testosterone also directly modulates dopamine activity within the striatum, and consistent with our findings, recent studies have shown that modulation of dopamine in the striatum can lead to changes in response bias (e.g. Rutledge et al, 2015; Rigoli et al, 2016). Overall, our results highlight the importance of a better understanding of the role of pubertal hormones in decision related processes and the development of a more nuanced view on the complexities of adolescent decision making (Casey, Galván, & Somerville, 2015).

The current findings also have implications that go beyond the domain of decision making, such as instrumental learning (Delgado, Miller, Inati, & Phelps, 2005; O’Doherty, 2004). Specifically, the dorsal striatum is involved in adaptively setting the balance between model-based versus model-free learning (Daw, Niv, & Dayan, 2005; Dayan & Berridge, 2014). Model-based learning entails having a complex model of the world, from which internal representations are used to guide goal-directed behavior. Engagement of the prefrontal cortex during model-based learning, specifically the dorsolateral prefrontal cortex and its coupling with the striatum ensures maintaining associations between actions and their respective future outcomes (Balleine & O’Doherty, 2010; Smittenaar, FitzGerald, Romei, Wright, & Dolan, 2013). In contrast, model-free learning is instead guided by experiences in a trial-and-error fashion, with the goal of solving a current task without requiring an understanding how the solution works. As such, adolescents’ drive to explore information in a way that is indifferent to the future utility of that information (Somerville et al., 2017).

The idea that testosterone in adolescence may modulate we could expect that testosterone may reduce the influence of the prefrontal cortex and thus tip the balance towards more model-free learning (see also Decker, Otto, Daw, & Hartley, 2016). This proposed mechanisms may explain recent findings in the literature, such as the increasing specific components of executive functioning is in line with a recent study by Nguyen et al. (2017) which demonstrated a negative relationship between baseline testosterone levels and reported performance on action monitoring and behavioral flexibility as measured by the Behavior Rating Inventory of Executive Function (BRIEF) in a longitudinal adolescent sample. Interestingly, this behavioral effect was related to a positive prefrontal-hippocampal covariance in structural grey matter development. Similarly, Piekarski, Boivin, & Wilbrecht (2017) showed that prepubertal hormone treatment in mice led them to require more trials to reach a criterion during a reversal-learning task, while maturation of tonic and phasic inhibition (a mechanisms of neuroplasticity) in the frontal cortex was accelerated. Related to learning, our findings have implications for our understanding of the interplay between testosterone and risk taking. Most of the time adolescents do not have full knowledge about probabilities and outcomes, but learn about these through experience (van den Bos & Hertwig, 2017). To mimic such a process, several studies have relied on the Balloon Analog Risk Task (BART; Lejuez et al., 2002), or similar tasks (Iowa Gambling Task, Driving Game, Sampling paradigm) to measure risk behavior (e.g., Aklin, Lejuez, Zvolensky, Kahler, & Gwadz, 2005; MacPherson, Magidson, Reynolds, Kahler, & Lejuez, 2010; Romer et al., 2009; Telzer, Fuligni, Lieberman, & Galván, 2013). Using this task, Peper and colleagues (2013) concluded that higher levels of testosterone were associated with higher levels of risk taking. On the basis of our findings it could be hypothesized that it is specifically the role of testosterone in instrumental learning that leads to differences in risk taking, instead of modulating risk preferences directly. In other words, instead of being risk-taking per se, adolescents may have difficulties learning action-outcome contingencies (in the BART, learning the association between number of pumps and the eventual burst of the balloon). Studies separating risk from learning aspects would deliver important insights into this proposed hypothesis.

In contrast with our previous study (Laube et al., 2017), we did not find an immediacy effect, nor did we find that magnitude of this effect was related to testosterone. Although the current task was not optimally designed to test for such an effect, this was nevertheless an unexpected result. One potential reason for the lack of an observable immediacy effect may be the use of monetary rewards only in the current study. Previous work suggests that the immediacy effect is strongest when individuals are presented with non-monetary rewards such as movie tickets (Read, Loewenstein, & Kalyanaraman, 1999), food choices (Read & Van Leeuwen, 1998) and primary rewards (McClure, Ericson, Laibson, Loewenstein, & Cohen, 2007), possibly because these rewards can be enjoyed immediately. Thus, future studies may seek to include both monetary and non-monetary rewards to distinguish immediacy effects from more general temporal discounting in adolescent populations.

Finally, there are several caveats in the current study. The current sample assessed male participants only. Although circulating testosterone is higher in males than in females, it also increases in females during puberty. Recent large-scale studies suggest gender differences in both impulsivity (Cross, Copping, & Campbell, 2011) and sensation seeking (D’Acremont & Van Der Linden, 2005; Steinberg et al., 2008). In addition, gonadal hormones have been associated with variation in the speed of brain maturation during adolescence (Raznahan et al., 2010). Future studies should include female adolescents to assess the role of pubertal testosterone increases in adolescent impatience in females. Second, longitudinal data is needed to fully distinguish pubertal maturation from non-developmental individual differences in testosterone levels. Although the relationship between testosterone levels and PDS suggests that testosterone is indeed capturing developmental effects, additional longitudinal investigation would increase our confidence about the conclusion that we report puberty-related effects. Third, although even the non-incentivized intertemporal choice task has a strong record in correlating with real-world outcomes such as earnings, education, and early death in adolescents (Bickel, Odum, & Madden, 1999; Golsteyn, Grönqvist, & Lindahl, 2014; Petry, 2001; Reimers, Maylor, Stewart, & Chater, 2009), the present data allow only indirect inferences about the relationship between real-world outcomes and pubertal testosterone.

In conclusion, the current study provides novel insights into the underlying mechanism of developmental changes in impulsive behavior across adolescence. To our knowledge, the current study is the first one to show an association between pubertal testosterone and choice related brain activation. By using a multimodal approach combining fMRI, hormonal assessment, and cognitive modeling of task-related behavior, we found support for the hypothesis that pubertal testosterone modulates the dorsal striatal pathway, which in turn may bias adolescents to impulsive behavior. Note that impulsive behavior is not harmful in itself, and it may also be even specifically beneficial in this developmental period (Spear, 2010, van den Bos, Laube & Hertwig, in press) Recent research suggests that pubertal hormones also regulate plasticity in the prefrontal cortex (Juraska & Willing, 2017; Piekarski et al., 2017). Thus, although pubertal changes in testosterone may increase impulsivity, these changes may also provide a unique window of opportunity to train self-control (see Berkman, Graham, & Fisher, 2012) and the social-cognitive skills (Crone & Dahl, 2012) necessary to successfully navigate in a social world.

