## Supplementary Material for "Pubertal Testosterone Correlates with Adolescent Impatience and Dorsal Striatal Activity"

**Supplementary Materials**

**Table S1**

*Descriptives of the testosterone and PDS measures*

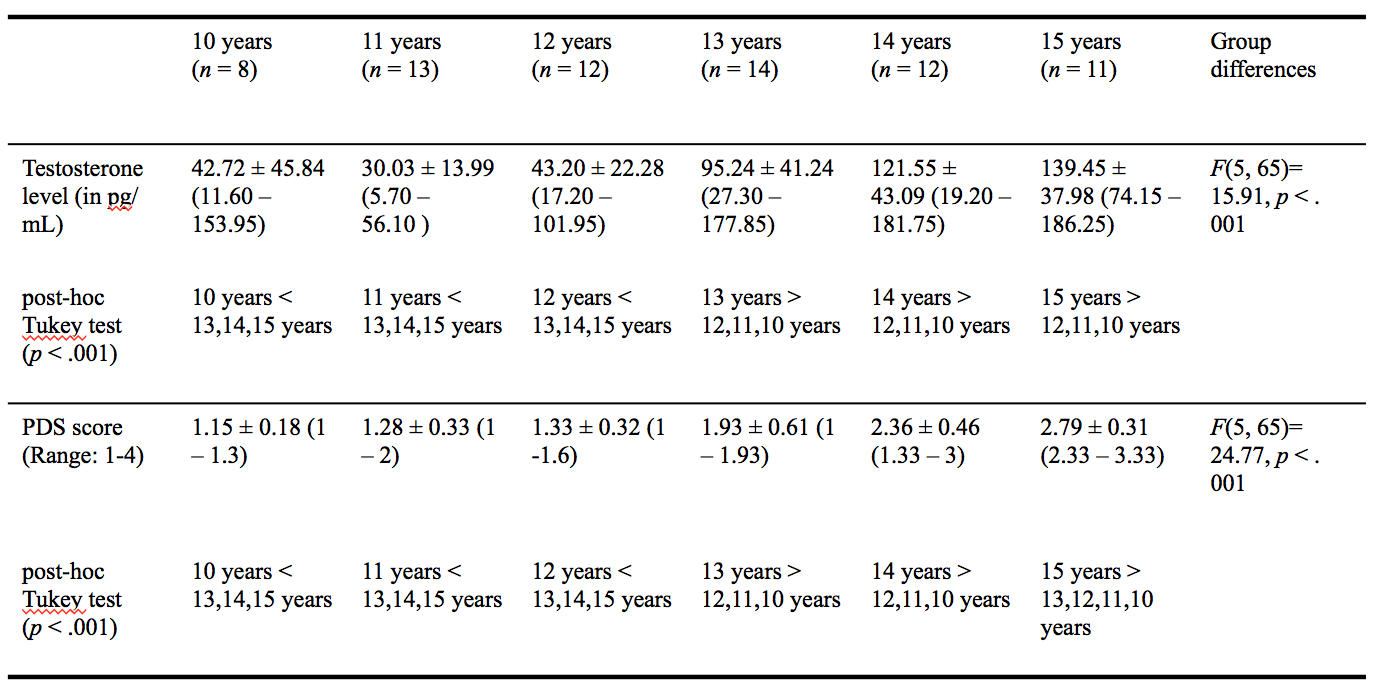

*Table S1: M* ± *SD* (and range) for testosterone and average PDS score for age groups 10, 11, 12, 13, 14 and 15 years, respectively. In addition, group differences are calculated based on log testosterone levels and average PDS scores.

PDS: Pubertal Developmental Scale

*N* = 70

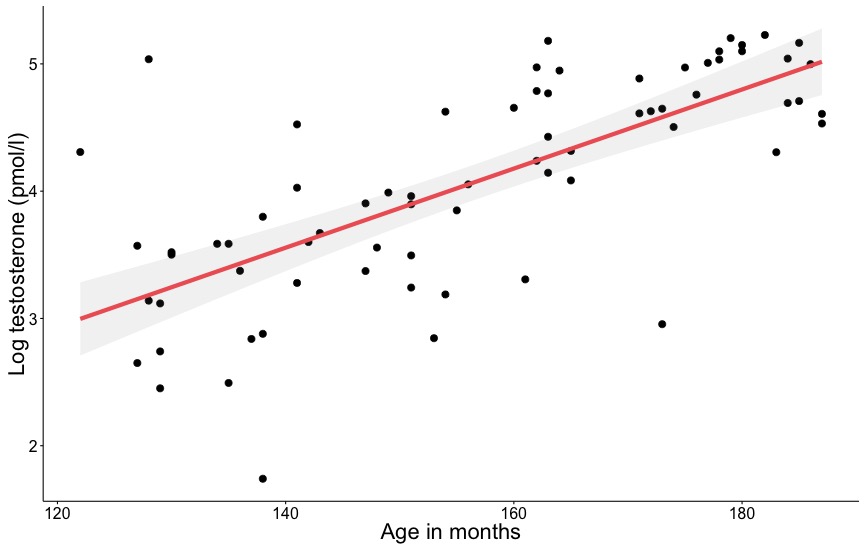

*Figure S1.* A graphic representation of the relationship between log testosterone levels and age (in months). Shaded area represents 95% confidence interval.

**Stimuli set for scanner session**

**Procedure.** Hypothetical monetary rewards were generated from an independent task prior to the MRI session. This task was part of a different study, which aimed at investigating the role of integral affect in intertemporal choice. First, participants chose between 30 leisure activities, where each activity was presented with an image printed on a paper card. Of the 30 options available, they chose the 12 cards they liked best and then ranked those 12 activities from most to least pleasant. Next, participants indicated their willingness-to-pay (WTP; in euro; Pachur et al., 2014; Rottenstreich & Hsee, 2001; Suter, Pachur, & Hertwig, 2015; Suter, Pachur, Hertwig, et al., 2015), To ensure that participants’ WTPs did not reflect actual prices in the real world, we chose outcomes that either were not common in everyday life (e.g., driving a Ferrari) or entailed no monetary costs (e.g., going to a lake with friends). Next, participants completed the adaptive intertemporal choice task (as described in the paper under 2.3). After the participants completed 60 trials on this task, we fitted data with the hyperbolic discount function which parameter estimates were used to generate the trials of the scanner session:

Eq. 1$V=\frac{A}{(1+kD)}$

where A is the amount in euros of the LL reward in the now/later condition, as well as the SS reward in the later/later condition. The individual discount rates that resulted from this procedure were used to generate the choice set for the final fMRI intertemporal choice paradigm. Specifically, we used the following logistic choice rule to predict choices by mapping the models predicted subjective value for the smaller sooner (*V_SS_* ) and larger later (*V_LL_* )options onto a probability of choosing the larger later (*P_LL_*) option. Here, we used four different probabilities (20, 40, 60 and 80 % for choosing the LL option) in order to manipulate choice conflict or difficulty.

Eq. 2 ${\text{ }\text{P}}_{\text{LL}}\text{=}\frac{\text{1}}{\text{1}+e^{-\theta(V_{LL}-V_{SS})}}$,

where $\theta$ estimates response noise. This function assumed that each individual would choose the option with the highest subjective value with the highest probability.

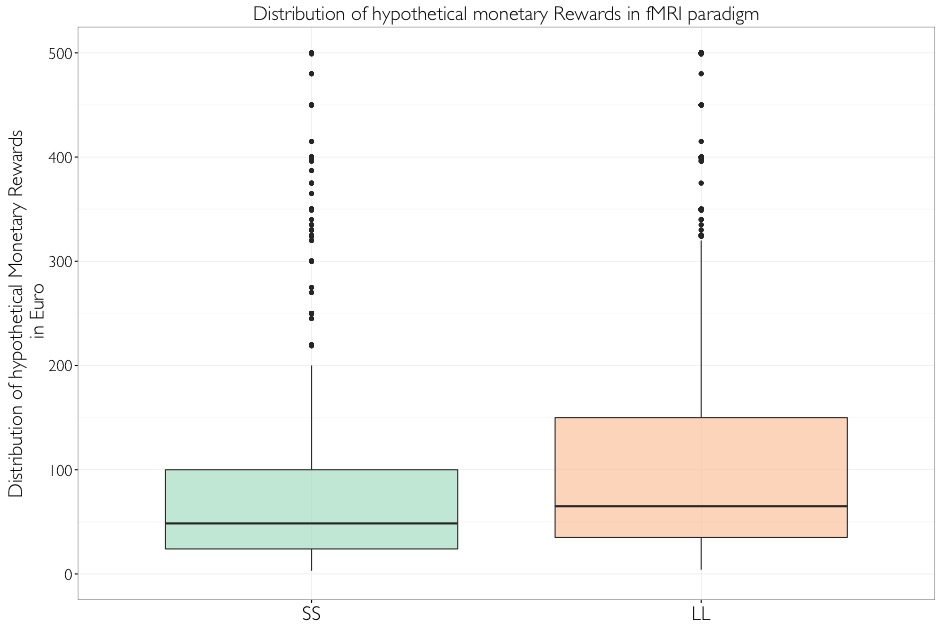

*Figure S2.* Distributions of monetary amounts used in study, separately for smaller sooner (SS) choices (left) and larger later (LL) choices (right).

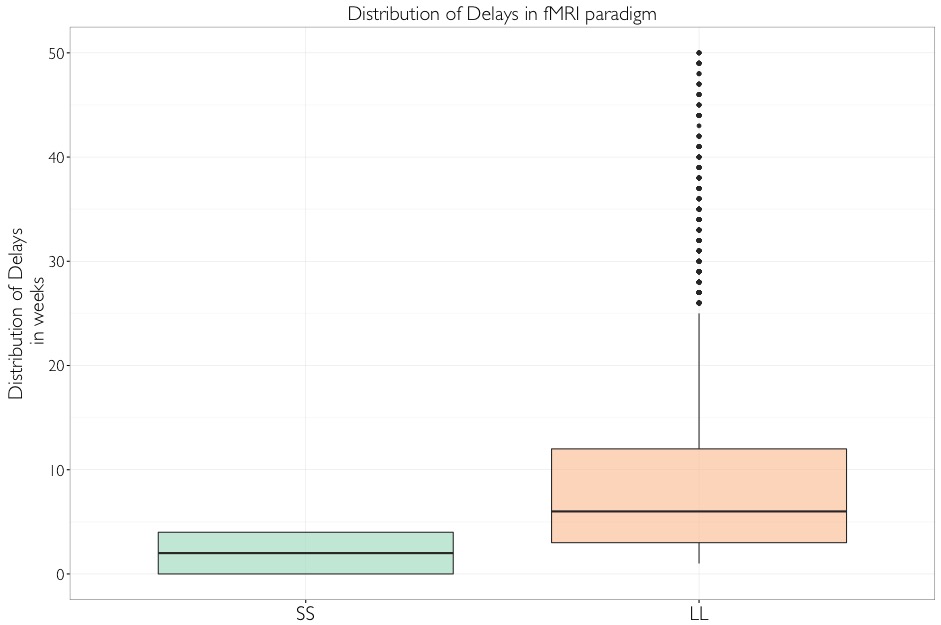

*Figure S3.* Distributions of delays used in study, separately for smaller sooner (SS) choices (left) and larger later (LL) choices (right).

**Table S2**

*Median of respective stimuli and its correlation with age, testosterone and PDS*

| Median | Variable | Age | Testosterone | PDS |
| --- | --- | --- | --- | --- |
| 55.41 | SS | .06 | -.08 | .04 |
| 75.22 | LL | .07 | -.08 | .04 |
| 9.27 | Delay | -.23 | -.03 | -.12 |

*Table S2:* Table showing the median for each type of stimuli (SS, LL and delay) used in scanner task, and their respective correlations with age, testosterone and PDS.

PDS: Pubertal Developmental Scale, *N* = 48

**Table S3**

| Mean  (SD) | Variable | Age | Testosterone | PDS | CFT |
| --- | --- | --- | --- | --- | --- |
| 158.66  (9.29) | Age (in months) | 1 |  |  |  |
| 70.77  (42.81) | Testosterone (pmol/l) | .52**  [.47, .56] | 1 |  |  |
| 1.70  (0.54) | PDS | .57***  [.53, .61] | .65***  [.61, .68] | 1 |  |
| 11.88  (2.24) | CFT | -.10  [-.16, -.04] | .18  [.12, .24] | -.17  [-.23, -.11] | 1 |

*Zero-order correlations between study variables for the sub group*

*Table S3:* Table showing the means and standard deviations for each of the study variables (left) and zero order correlations along with 95% confidence intervals (right) for the subsample including *N* = 32 participants.

PDS: Pubertal Developmental Scale, CFT: Culture Fair Test

****p* < .001, ***p* < .01, **p* < .05., *N*=32

**Table S4**

| Mean  (SD) | Variable | Age | Testosterone | PDS | CFT |
| --- | --- | --- | --- | --- | --- |
| 156.10  (19.90) | Age (in months) | 1 |  |  |  |
| 75.19  (55.13) | Testosterone (pmol/l) | .67***  [.63, .70] | 1 |  |  |
| 1.82  (0.71) | PDS | .82***  [.80, .84] | .72***  [.69, .75] | 1 |  |
| 11.11  (2.84) | CFT | .26  [.20, .32] | .02  [-.04, .08] | .12  [.06, .18] | 1 |

*Zero-order correlations between study variables for the imaging group*

*Table S4:* Table showing the means and standard deviations for each of the study variables (left) and zero order correlations along with 95% confidence intervals (right) for the imaging sample including *N* = 48 participants.

PDS: Pubertal Developmental Scale, CFT: Culture Fair Test

****p* < .001, ***p* < .01, **p* < .05., *N*=48

**Imaging Analysis with sub groups**

**NAcc and testosterone.** We did not find a significant relationship between testosterone levels and NAcc activity for LL over SS choices for neither the total group (*N* = 70), *b* = -.28, 95% CI (-.62, .06), *p* = .11, nor the subgroup (*N* = 32), *b* = -.17, 95% CI (-.61, .27), *p* = .44.

**Caudate and testosterone.** We did find a significant relationship between testosterone levels and caudate activity for the difference between SS and LL choices in both the whole group (*N* = 70), *b* = -.48, 95% CI (-.80, -.16), *p* < .005, as well as in the subgroup (*N* = 32), *b* = -.52, 95% CI (-.92, -.12), *p* = .01.

**Activity in caudate related to testosterone and bias parameter.** In the whole group (*N* = 70), we found a positive relationship between activation within the caudate nucleus for SS over LL choices and the bias parameter, *b* = .04, 95% CI (-.21, .29), *p* = .75, but results were also not statistically significant. For the subgroup (*N* = 32), the relationship was negative, but also not statistically significant, *b* = -.31, 95% CI (-.79, .16), *p* = .09.
